# Tumor evolution of glioma intrinsic gene expression subtype associates with immunological changes in the microenvironment

**DOI:** 10.1101/052076

**Authors:** Qianghu Wang, Xin Hu, Baoli Hu, Florian Muller, Hoon Kim, Massimo Squatrito, Tom Millelsen, Lisa Scarpace, Floris Barthel, Yu-Hsi Lin, Nikunj Satani, Emmanuel Martinez-Ledesma, Edward Chang, Adriana Olar, Guocan Wang, Ana C. deCarvalho, Eskil Eskilsson, Siyuan Zheng, Amy B. Heimberger, Erik P. Sulman, Do-Hyun Nam, Roel G.W. Verhaak

**Affiliations:** Department of Genomic Medicine, The University of Texas MD Anderson Cancer Center, Houston, TX 77030, USA; Department of Radiation Oncology, The University of Texas MD Anderson Cancer Center, Houston, TX 77030, USA; Department of Bioinformatics and Computational Biology, The University of Texas MD Anderson Cancer Center, Houston, TX 77030, USA; Department of Cancer Biology, The University of Texas MD Anderson Cancer Center, Houston, TX 77030, USA; Department of Cancer Systems Imaging, The University of Texas MD Anderson Cancer Center, Houston, TX 77030, USA; Department of Pathology, The University of Texas MD Anderson Cancer Center, Houston, TX 77030, USA; Department of Neurosurgery, The University of Texas MD Anderson Cancer Center, Houston, TX 77030, USA; University of Texas-Houston Graduate School in Biomedical Sciences, Houston, TX 77030, USA; Cancer Cell Biology Programme, Seve Ballesteros Foundation Brain Tumor Group, Centro Nacional de Investigaciones Oncológicas, CNIO, 28029 Madrid, Spain; Departments of Neurology and Neurosurgery, Henry Ford Hospital, Detroit, MI 48202, USA; Oncology Graduate School Amsterdam, VU University Medical Center, 1081 HV Amsterdam, The Netherlands; Institute for Refractory Cancer Research, Samsung Medical Center, Seoul 06351, Korea; Department of Health Sciences and Technology, Samsung Advanced Institute for Health Sciences and Technology, Sungkyunkwan University, Seoul 06351, Korea; Department of Neurosurgery Samsung Medical Center, Sungkyunkwan University School of Medicine, Seoul, 135-710, Korea

**Keywords:** glioblastoma, disease recurrence, mesenchymal subtype, proneural to mesenchymal transition, gene expression profiling, tumor microenvironment, macrophages/microglia, immune cells

## Abstract

**Summary:** We leveraged IDH wild type glioblastomas and derivative neurospheres to define tumor-intrinsic transcription phenotypes. Transcriptomic multiplicity correlated with increased intratumoral heterogeneity and tumor microenvironment presence. In silico cell sorting demonstrated that M2 macrophages/microglia are the most frequent type of immune cells in the glioma microenvironment, followed by CD4 T lymphocytes and neutrophils. Hypermutation associated with CD8+ T cell enrichment. Longitudinal transcriptome analysis of 124 pairs of primary and recurrent gliomas showed expression subtype is retained in 53% of cases with no proneural to mesenchymal transition being apparent. Inference of the tumor microenvironment through gene signatures revealed a decrease in invading monocytes but a subtype dependent increase in M2 macrophages/microglia cells after disease recurrence. All expression datasets are accessible through http://recur.bioinfo.cnio.es/.

**Significance:** IDH wild type glioblastoma expression phenotypes have been related to tumor characteristics including genomic abnormalities and treatment response. We explored the intratumoral transcriptomic landscape, including a definition of tumor-intrinsic gene expression subtypes and how they relate to the different cellular components of the tumor immune environment. Comparison of matching primary and recurrent gliomas provided insights into the treatment-induced phenotypic tumor evolution. Proneural to mesenchymal transitions have long been suspected but were not apparent, while intratumoral heterogeneity was a predictor of subtype transition upon recurrence. Characterizing the evolving glioblastoma transcriptome en tumor microenvironment aids in designing more effective immunotherapy trials. Our study provides a comprehensive transcriptional and cellular landscape of IDH wild type GBM during treatment modulated tumor evolution.

**Highlights:**

- Next generation GBM-intrinsic transcriptional subtypes: proneural, classical, mesenchymal
- M2 macrophages, CD4+ T-lymphocytes and neutrophils dominate glioblastoma microenvironment
- Sensitivity to radiotherapy may associate with M2 macrophage presence
- CD8+ T cells are enriched in hypermutated GBMs at diagnosis and recurrence

## INTRODUCTION

The intrinsic capacity of glioblastoma (GBM) tumor cells to infiltrate normal brain impedes surgical eradication and predictably results in high rates of early recurrence. To better understand determinants of GBM tumor evolution and treatment resistance, The Cancer Genome Atlas Consortium (TCGA) performed high dimensional profiling and molecular classification of nearly 600 GBM tumors (Brennan et al., 2013; Cancer Genome Atlas Research, 2008; Ceccarelli et al., 2016; Noushmehr et al., 2010; Verhaak et al., 2010b). In addition to revealing common mutations in genes such as *TP53*, *EGFR*, *IDH1*, and *PTEN*, as well as the frequent and concurrent presence of abnormalities in the p53, RB and receptor tyrosine kinase pathways. Unsupervised transcriptome analysis identified four clusters, referred to as classical, mesenchymal, neural and proneural, that were tightly associated with genomic abnormalities(Verhaak et al., 2010a). The proneural and the mesenchymal expression subtypes have been most consistently described in literature with proneural relating to a more favorable outcome and mesenchymal to unfavorable survival (Huse et al., 2011; Phillips et al., 2006; Zheng et al., 2012), but these findings were affected by the relatively favorable outcome of IDH-mutant glioblastoma which are consistently classified as proneural (Noushmehr et al., 2010; Verhaak et al., 2010a). Proneural to mesenchymal switching upon disease recurrence has been described as a source for treatment resistance in GBM relapse (Bao et al., 2006; Bhat et al., 2013; Ozawa et al., 2014; Phillips et al., 2006), but the relevance of this phenomenon in glioma progress remains ambiguous.

GBM tumor cells along with the tumor microenvironment create a complex milieu that ultimately promotes tumor cell plasticity and disease progression (Olar and Aldape, 2014). The presence of tumor-associated stroma results in a mesenchymal tumor gene signature and poor prognosis in colon cancers (Isella et al., 2015). Furthermore, the association between a mesenchymal gene expression signature and reduced tumor purity has been identified as a common theme across cancer (Martinez et al., 2015; Yoshihara et al., 2013). Tumor-associated macrophages/microglia in GBM have been proposed as regulators of proneural-to-mesenchymal transition through NF-kB activation (Bhat et al., 2013) and may provide growth factor mediated proliferative signals, which could be therapeutically targeted (Patel et al., 2014; Pyonteck et al., 2013; Yan et al., 2015).

Here, we explored the properties of the microenvironment in different GBM gene expression subtypes and characterized the transition between molecular subtypes before and after therapeutic intervention. In doing so, we improved the robustness of gene expression subtype classification through revised gene signatures and proposed analytical methodology. Our results suggested that the tumor microenvironment interferes with expression based classification of GBM, both at the primary disease stage as well as at disease recurrence, and suggest a role for the macrophage/microglia in treatment response.

## RESULTS

### Harnessing glioma sphere-forming cells identifies GBM specific intertumoral transcriptional heterogeneity

We set out to elucidate the tumor-intrinsic and tumor microenvironment independent transcriptional heterogeneity of GBMs. We performed a pairwise gene expression comparison of independent set of GBMs and the derivative glioma sphere-forming cells (GSCs) (n = 37) (Galli et al., 2004). In total, 5,334 genes were found to be significantly higher expressed in parental GBMs relative to derived GSCs that could be attributed by the tumor associated GBM microenvironment (Figure 1A). To focus the analysis on the tumor-intrinsic transcriptome, these genes were filtered from further analysis. GBMs with IDH mutations have distinct biological properties and favorable clinical outcomes compared to IDH wild-type GBMs (Brennan et al., 2013; Cancer Genome Atlas Research et al., 2015; Ceccarelli et al., 2016; Noushmehr et al., 2010). Using the filtered gene set, we performed consensus non-negative matrix factorization clustering to identify three distinct subgroups amongst 369 IDH wild type GBMs (Figure 1B; Figure 1C). When comparing the clustering result with the previously defined proneural (PN), neural (NE), classical (CL) and mesenchymal (MES) classification (Brennan et al., 2013; Verhaak et al., 2010b), three subgroups were strongly enriched for CL, MES and PN GBMs, respectively (Figure S1). Consequently, we labeled the groups as CL, MES and PN. None of the three subgroups was enriched for the NE class, suggesting its neural phenotype is non-tumor specific. The NE group has previously been related to the tumor margin where normal neural tissue is more likely to be present (Gill et al., 2014; Sturm et al., 2012) and such contamination might explain why the neural subtype was the only subtype to lack characteristic gene abnormalities (Brennan et al., 2013; Verhaak et al., 2013). To be able to classify external GBM samples, we implemented a single sample gene set enrichment analysis (ssGSEA) based equivalent distribution resampling classification strategy using 70-gene signatures for each subgroup (**Table S1**)(Figure 1D), to assign each sample three empirical classification p-values by which we determined the significantly activated subtype(s) in the samples. We prepared an R-library to facilitate others to evaluate our approach (Supplementary File 1). Using this method we found that the stability of cluster assignments of 144 TCGA GBM samples profiled using both RNA sequencing and Affymetrix U133A microarrays was 95% concordance (Figure S2, **Table S2**). This was an improvement over the 77% subtype concordance determined using previously reported methods (Verhaak et al., 2010b). We evaluated the distribution of somatic variants across all the three molecular subtypes (Figure 1E)(Figure S3) and confirmed the strong associations between subtypes and genomic abnormalities in previously reported driver genes (Brennan et al., 2013; Verhaak et al., 2010b).

**Figure 1.**
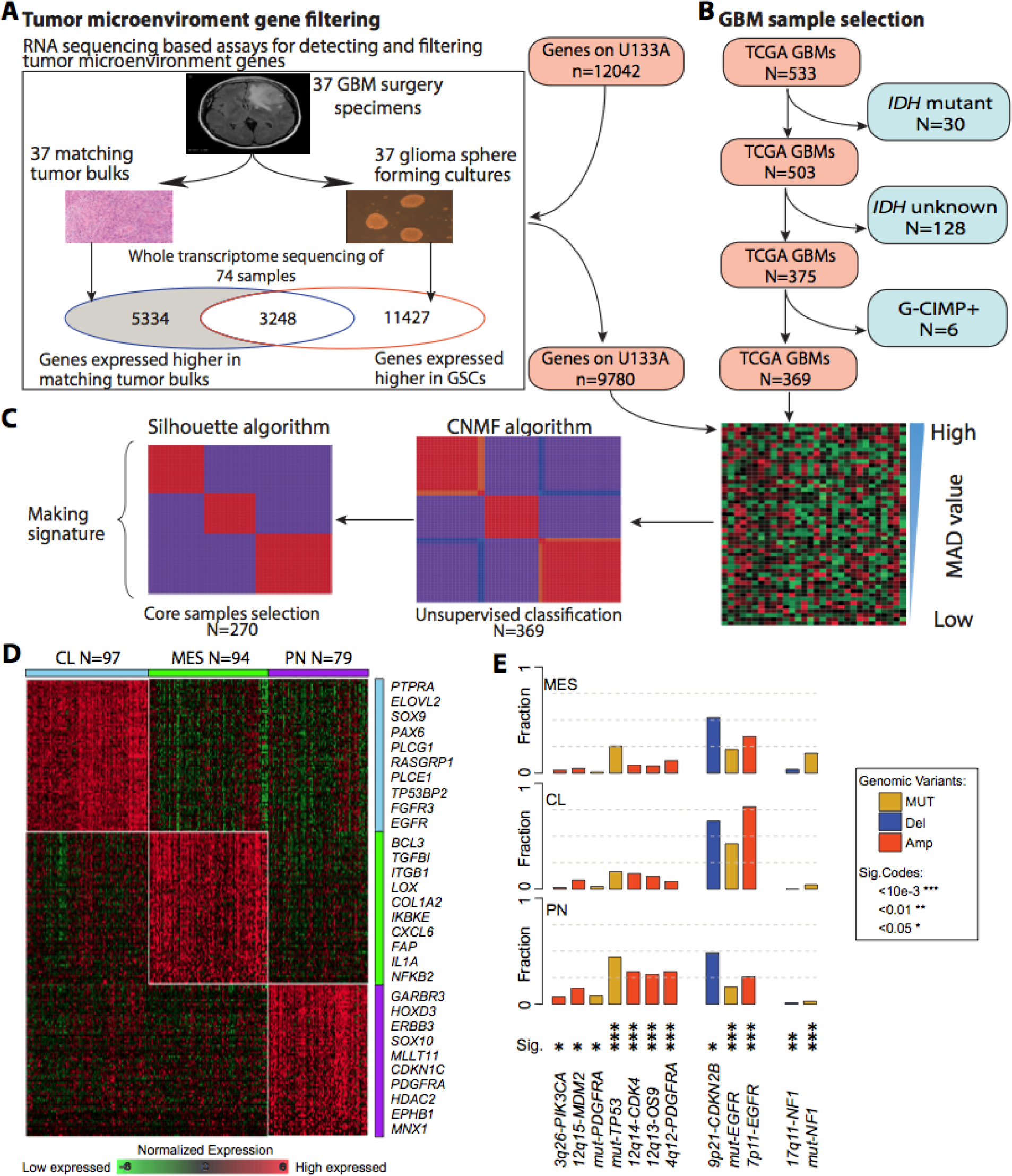
Molecular classification of IDHwt GBMs. **(A)** Filtering tumor associated microenvironment genes. (**B**) Discarding IDH mutation related GBMs. (**C**) Overview of NMF clustering. (**D**) Heatmap of 70-gene signatures by gene expression subtype were developed based on 270 GBMs. Top ten genes are shown for each subtype. (**E**) Frequency of subtype related somatic genomic alterations. Chi-square test was used to calculate the distribution difference among three subtypes per genomic variant.

### Multi-activation of subtype signatures associated with intratumoral heterogeneity

We observed that 34/369 (9.2%) samples showed significant enrichment of multiple ssGSEA scores (empirical classification p-value<0.05), suggesting these cases activate more than one transcriptional subtype (Figure 2A). To quantify this phenomenon, a score ranging from 0 to 1 was defined to quantitatively evaluate the simplicity of subtype activation based on order statistics of ssGSEA score. Samples with high simplicity scores activated a single subtype and those with lowest simplicity scores activated multiple subtypes. All multi-subtype TCGA samples showed simplicity scores of less than 0.1 (Figure 2A). To determine whether transcriptional heterogeneity associated with genomic intratumoral heterogeneity, we correlated simplicity scores, total mutation rates and subclonal mutation rates. Included in the analysis were 224 TCGA GBMs with available whole exome sequencing data (Kim et al., 2015) and ABSOLUTE (Carter et al., 2012) determined high tumor purity (> 0.8) to equalize the mutation detection sensitivity (Aran et al., 2015). Although not significant (Wilcoxon rank test p-value=0.143), the total mutation rate was less in the bottom 30% with lowest simplicity scores versus the top 30% samples with highest simplicity scores. The subclonal mutation rate was significantly higher (p-value=0.024) in samples with lowest simplicity scores (Figure 2B; **Table S3**), suggesting that increased intratumoral heterogeneity associates with increased transcriptional heterogeneity.

**Figure 2.**
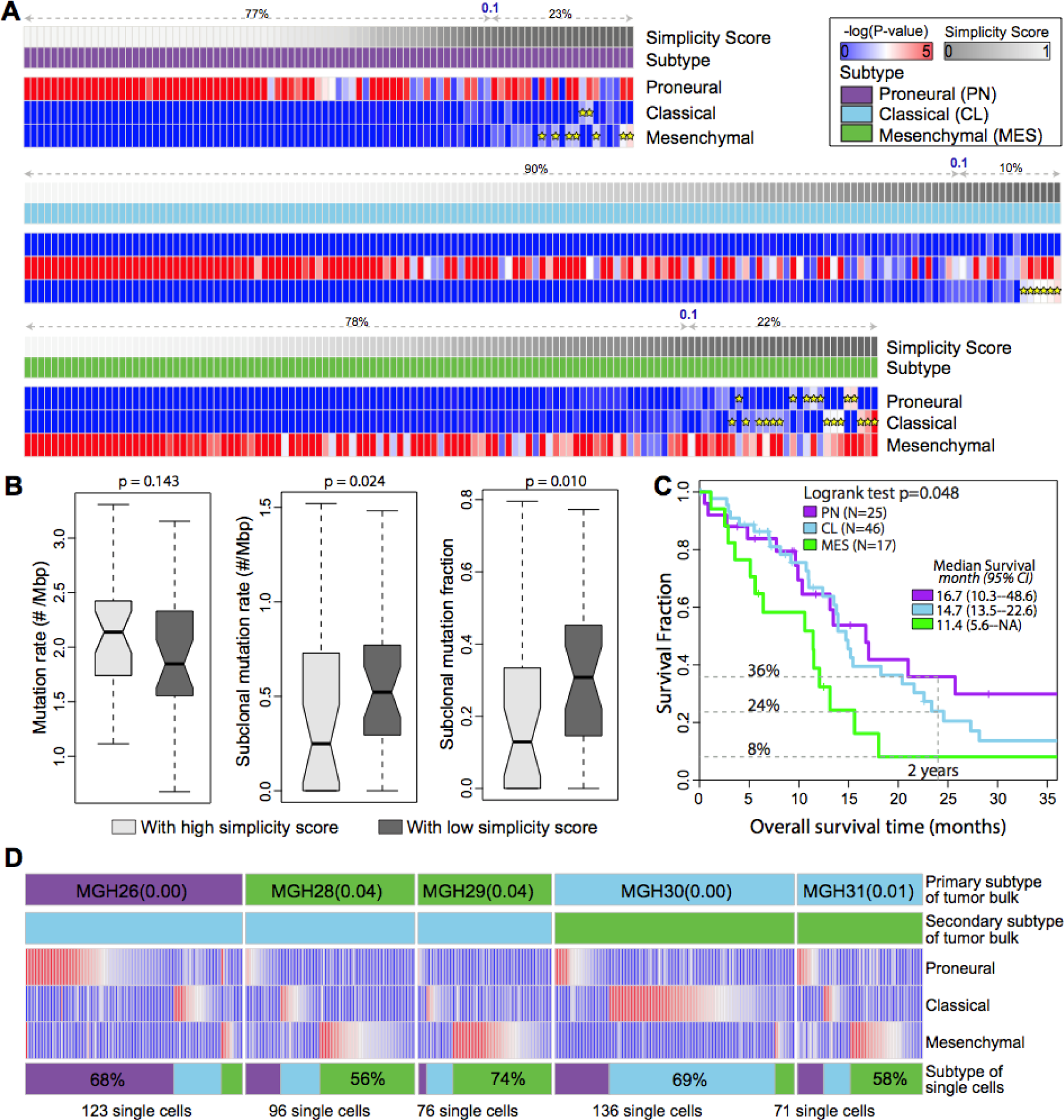
Multi activation of transcriptional subtypes associated with intratumoral heterogeneity. (**A**) The expression profiles of 369 IDHwt GBMs were analyzed using Affymetrix U133A. The empirical -log(empirical P-value) of raw ssGSEA enrichment scores at each signature are shown as heatmaps, with dark blue representing no activation and bright red as highly activated. Yellow star indicates the secondary activated subtype (empirical p-value<0.05). For each panel, the first row shows simplicity score, and the second row indicates transcriptional subtype. (**B**) Comparison of mutation rate, subclonal mutation rate and subclonal mutation fraction between IDHwt GBMs with high and low simplicity scores. P-values were calculated using Wilcoxon rank test and shown at the top of each panel. (**C**) Kaplan-Meier survival curve by subtype. (**D**) Transcriptome classification of five bulk tumor samples and 502 single GBM cells derived from them. The top two row of each panel show the dominant and secondary subtype of the GBM tumor bulk. The heatmap of each panel shows the empirical -log(P-value) of the ssGSEA scores of the derived single GBM cells on each of the three subtype signatures. The bottom row shows the subtype distribution of derived single GBM cells within the same GBM tumor of origin.

We compared outcomes amongst the three transcriptional groups and observed no significant differences (Figure S4). However, when restricting the analysis to samples with high simplicity scores, a clear trend of MES showing worst survival and PN the most favorable outcome became visible. For example, Kaplan-Meier analysis of 88 samples with simplicity scores >0.99 showed a median survival of 11.4, 14.7 and 16.7 months were detected in MES, CL and PN, respectively, which was significantly different (log rank test, p=0.048)(Figure 2C)(Figure S4)**(Table S4)**. Higher simplicity scores correlated with relative favorable outcome within the PN set, non-significant in the CL subtype, and correlated with relatively unfavorable survival in the MES class (Figure S5).

**Figure 3.**
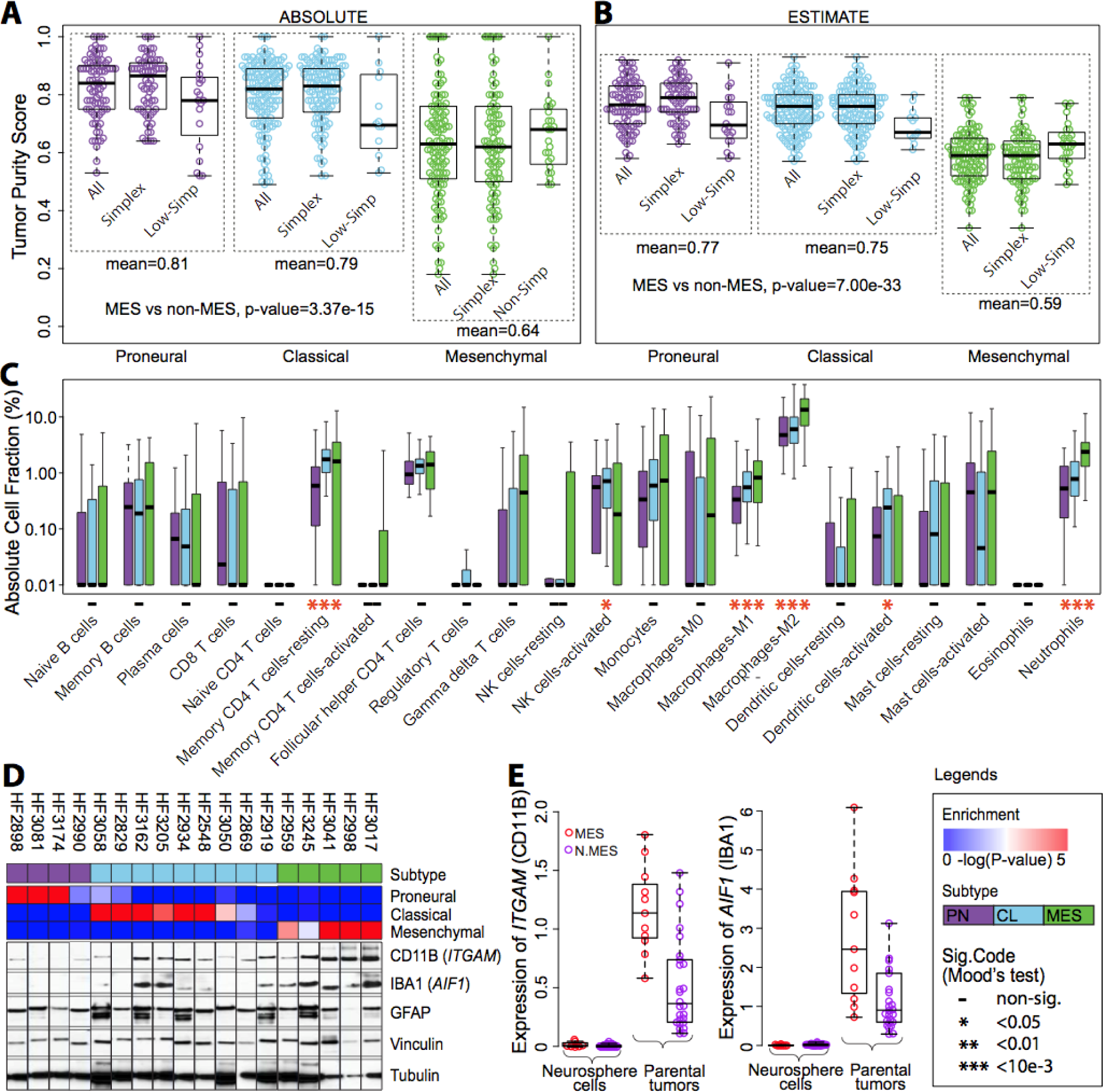
Transcriptional subtypes differentially activate the immune microenvironment. (**A, B**) Tumor purity of 364 respectively 369 TCGA IDHwt GBM samples was determined by ABSOLUTE and ESTIMATE. The difference in tumor purity between subtypes was evaluated using a two-sample heteroscedastic t-test. (**C**) Comparison of immune cell fractions among subtypes. Immune cell fractions were estimated using CIBERSORT and corrected using ABSOLUTE purity scores per sample. The distribution of immune cell fractions of 69 PN, 137 CL and 96 MES IDHwt GBMs with simplicity score>0.1 were shown by purple, skyblue and green boxplots, respectively. Median value difference of cell fraction among subtypes was evaluated using Mood’s test. (**D**) The upper panel shows ssGSEA enrichment scores and associated expression subtype classifications. Bottom panels display protein expression of the microglial markers integrin alpha M (ITGAM) and allograft inflammatory factor 1 (IBA1), astrocyte marker glial fibrillary acidic protein (GFAP) and the loading control tubulin. (**E**) Comparison of *ITGAM* and *IBA1* gene expression levels between GBM and derived neurosphere models.

Single GBM cell RNA sequencing recently suggested that GBMs are comprised of a mixture of tumor cells with variable GBM subtype footprints (Patel et al., 2014). We used this data to classify 502 single GBM cells in addition to the bulk tumor derived from five primary glioblastomas (**Table S5**). All bulk tumor samples showed simplicity scores less than 0.05 suggesting high transcriptional heterogeneity compared to 45 of 369 TCGA GBM samples with simplicity scores below 0.05 (Figure 2D). In four of five cases (MGH26, MGH28, MGH29 and MGH30), the bulk tumor samples were classified in the same primary subtype as the majority of their single cells (Figure 2D). Our analysis suggests that the heterogeneity observed at the single cell level is captured in the expression profile of the bulk tumor, and that the five GBM samples studied at the single cell level represented samples with relatively high transcriptional heterogeneity.

**Figure 4.**
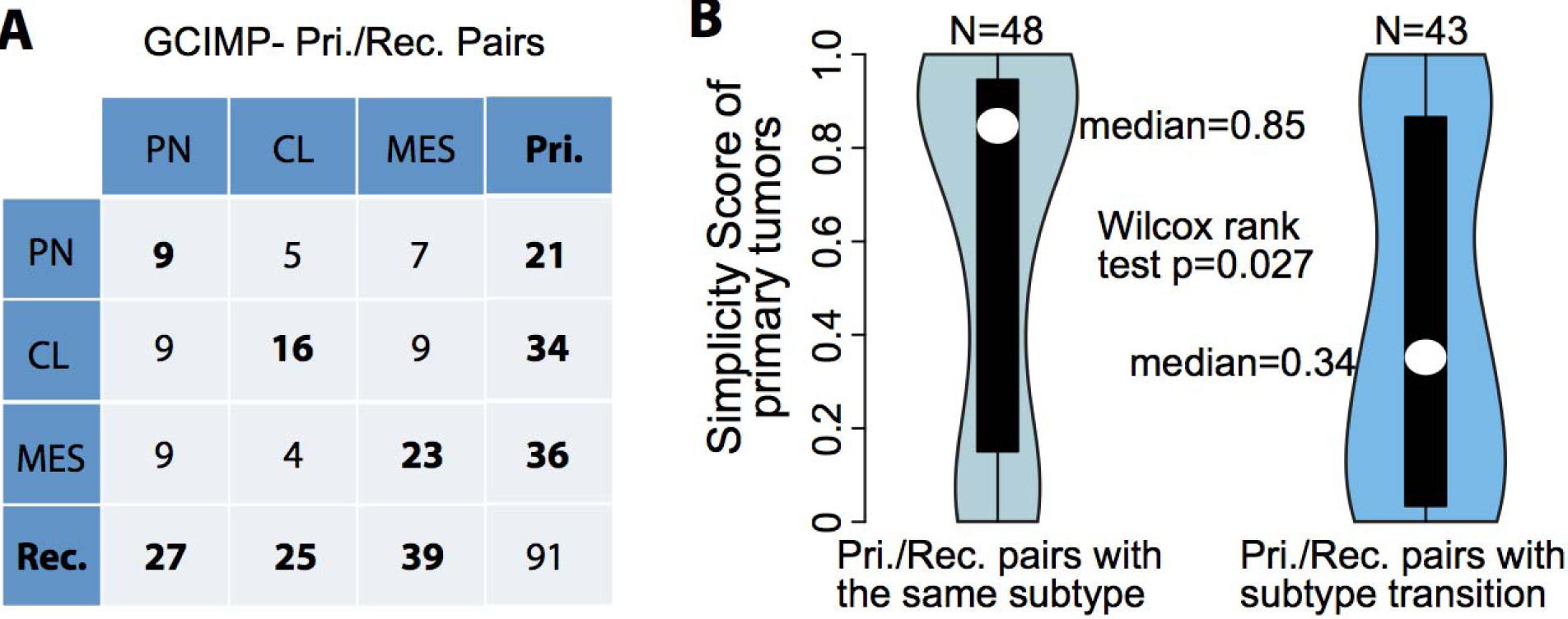
Comparison between transcriptional subtype of primary and paired recurrent tumors. (**A**) Rows and columns of the cross table represents subtype distribution frequency of primary and paired recurrent tumors, respectively. (**B**) Violin plots show the distribution of simplicity scores of pairs with (left) and without (right) subtype transition.

### Transcriptional subtypes differentially activate the immune microenvironment

Despite restricting the cluster analysis to genes exclusively expressed by GBM cells, we found that tumor purity predictions based on ABSOLUTE were significantly reduced in GBM classified as MES (Student T-test p-value < 10e-14; Figure 3A). This was corroborated by gene expression based predictions of tumor purity using the ESTIMATE method (Student T-test p-value < 10e-32; Figure 3B)(Yoshihara et al., 2013). The ESTIMATE method has been optimized to quantify tumor-associated fibroblasts and immune cells (Yoshihara et al., 2013) and the convergence of a decreased ABSOLUTE and decreased ESTIMATE tumor purity confirmed previous suggestions on the increased presence of microglial and neuroglial cells mesenchymal GBM (Bao et al., 2006; Engler et al., 2012; Gabrusiewicz et al., 2016; Ye et al., 2012). The mean simplicity score of samples classified as MES was 0.53 which was significantly lower than in PN (Wilcoxon rank test p-value<0.019) and CL subtypes (Wilcoxon rank test p-value<0.0001), confirming increased transcriptional heterogeneity. In order to identify genomic determinants of macrophage/microglia chemoattraction, we compared genomic alterations between mesenchymal class samples with high (n=51) and low (n=51) ABSOLUTE based tumor purity. GBM carrying hemyzygous loss of *NF1* or somatic mutations in *NF1* showed reduced tumor purity compared to GBM with wild type *NF1* (Wilcoxon rank test p-value=0.0007) and this association was similarly detected when limiting the analysis to MES samples (Wilcoxon rank test p-value=0.017)(Figure S6). Formation of dermal neurofibromas in the context of *Nf1* loss of heterozygosity has been reported to be context and microenvironment dependent (Le et al., 2009). Functional studies may clarify whether NF1 deficient GBM are able to recruit cells that provide them with a proliferative advantage, or whether *NF1* loss provides that proliferative advantage in a specific tumor-associated microenvironment context.

**Figure 5.**
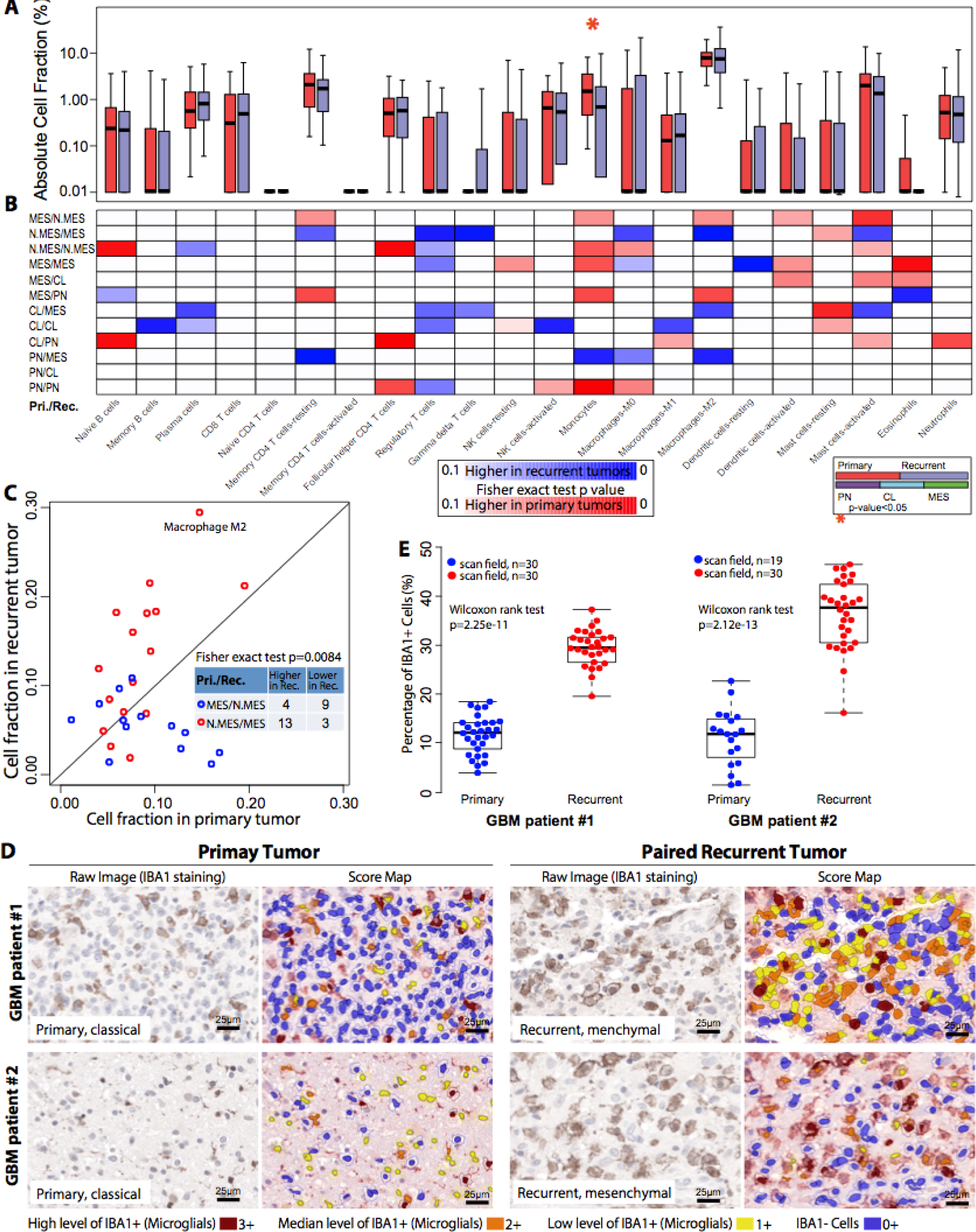
Microenvironment transition between 91 primary and paired recurrent IDH wild type GBM. (**A**) Red and blue boxplots represent the immune cell fraction distribution of each immune cell type. Immune cell fraction was calculated using CIBERSORT and adjusted using ESTIMATE purity scores. Difference between cell fraction of primary and paired recurrent tumors was calculated using Wilcoxon rank test. (**B**) The blue-to-red heatmap represents immune cell fraction changes upon tumor recurrence per subtype transitions which were list on the left of the heatmap. Fisher exact test was used to evaluate the distribution difference between patients with higher/lower immune cell fractions at tumor recurrence per subtype transition. (**C**) Each dot represents a pair of primary and recurrent GBM with axes indicating M2 macrophage cell fraction. (**D**) Representative images of IBA1 immunohistochemical staining and corresponding score map obtained by InForm image analysis in two matched pairs of primary and recurrent GBM. Scale bar, 25 µm. **(E)** Unbiased quantification of IBA1+ percentage in primary and recurrent GBMs.

To determine the cellular components of the tumor microenvironment across different transcriptional subtypes, we used the CIBERSORT in silico cytometry (Newman et al., 2015) method to evaluate absolute immune cell fractions. We evaluated 22 different immune cell types in 69 PN, 137 CL and 96 MES samples, after filtering samples with classification simplicity scores less than 0.1 (**Table S6**). Microglia are the resident macrophages in the central nervous system. Peripheral blood monocytes also give rise to tumor associated macrophages. These innate immune cells can be broadly classified as the proinflammatory M1 type and the alternative tumor promoting M2 type(Hambardzumyan et al., 2015). The M2 macrophage gene signature showed a greater association with the MES subtype (13.4%) relative to the PN (4.6%) and CL (6.0%)(Figure 3C), consistent with previous analysis of the TCGA database(Doucette et al., 2013; Gabrusiewicz et al., 2016). In addition to the M2 macrophage gene signature, there was also a significantly higher fraction of MES samples that expressed M1 macrophage (Student T-test p-value 3.20E-5) and neutrophil (Student T-test p-value 1.30E-9) gene signatures. In contrast, the activated natural killer T-cell gene signature (Student T-test p-value 4.91E-2) was significantly reduced in the MES subtype and resting memory CD4+ T cells (Student T-test p-value 5.40E-7) were less frequently expressed in the PN subtype.

To confirm the association of macrophages/microglia with the MES GBM subtype, we assessed protein expression levels of the ITGAM (alternatively known as CD11B) and IBA1 (also known as AIF1) macrophage/microglial markers in a set of 18 GBM for which we also characterized the expression subtype (Figure 3D) as well as through immunohistochemistry (Figure S7). We confirmed the microenvironment as the main source for *ITGAM/IBA1* transcription by comparing transcriptional levels in 37 GBM-neurosphere pairs used for gene filtering, which showed that neurospheres do not express *ITGAM/IBA1* (Figure 3E). The association of the MES GBM subtype with increased level of M2 microglia/macrophages may suggest that in particular MES GBM are candidates for therapies directed against tumor-associated macrophages(Pyonteck et al., 2013). Activated dendritic cell signatures (Student T-test p-value 7.36E-3)(Figure 3C) were significantly higher in the CL subtype, suggesting this subtype may benefit from dendritic cell vaccines (Palucka and Banchereau, 2012). Dendritic cells may require an activated phenotype in order to direct the immune system. A previous study suggested that MES GBM patients treated with dendritic cells were more likely to benefit (Prins et al., 2011).

**Figure 6.**
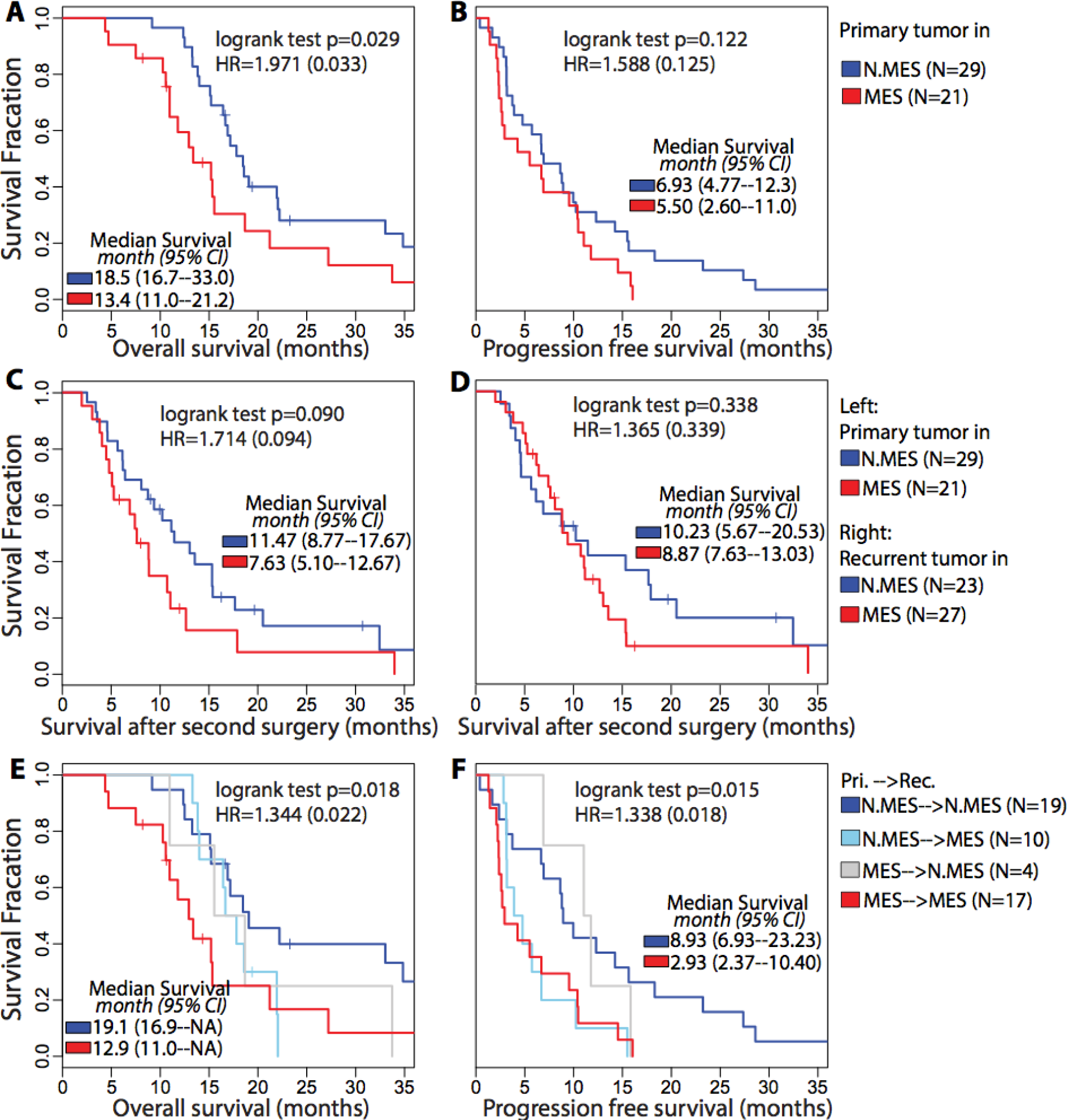
Survival analysis of paired IDH wild type GBM. (**A, B**) OS and PFS analyses between samples with different primary subtype. (**C**) Difference of survival time after secondary surgery between patients with non-MES and MES in primary tumors. (**D**) Survival analysis of time after secondary surgery between patients with non-MES and MES in recurrent tumors. (E, F) OS and PFS analyses between samples with difference recurrent subtype.

### Phenotypic plasticity upon GBM recurrence

Glioblastoma has long been hypothesized to progress along a proneural to mesenchymal axis(Phillips et al., 2006). To determine the relevance of this transition process in IDH wildtype glioma evolution, we performed a longitudinal analysis of the subtype classification and tumor-associated microenvironment in sample pairs obtained at diagnosis and first disease recurrence from 124 glioma patients. The cohort included 96 initial GBM and first recurrence, eight pairs of primary low grade glioma and matching secondary GBM, and 20 pairs of primary and recurrent low grade glioma. Gene expression profiles of 78 tumor pairs were analyzed through transcriptome sequencing, and remaining pairs were generated using Affymetrix (n = 31) and Illumina (n = 15) microarray, respectively. To facilitate exploration of this dataset we have made it available through the GlioVis portal http://recur.bioinfo.cnio.es/. We used a gene expression signature(Baysan et al., 2012) to determine that 33 of 124 cases were IDHmutant/GCIMP at presentation and recurrence (**Table S7**). We used the renewed gene signatures and classification method to determine molecular subtype of the 91 pairs of IDH wild type cases and found that expression class remained consistent after disease recurrence for 48 of 91 IDH-wildtype cases (52%)(Figure 4A). The MES subtype was most stable (64%) while the CL (47%) and PN (43%) phenotypes were less frequently retained. Nine, sixteen and eighteen post-treatment tumors switched subtypes to become CL, MES and PN at disease recurrence, respectively, indicating that PN and MES increased in frequency after recurrence while the CL subtype was least frequently found (Figure 4A). The CL expression class was previously found to be most sensitive to intensive therapy and it is possible that therapy provides a competitive advantage for non-CL cells, which could explain the reduced post-treatment incidence(Verhaak et al., 2010b). Our results did not identify enrichment for proneural to mesenchymal transitions.

We observed a significant difference in transcriptional simplicity between primary GBM retaining their expression class, versus those that switched to a different phenotype (Figure 4B). GBMs with a primary tumor simplicity score greater than 0.5, indicated lower transcriptional heterogeneity, were classified as the same subtype in 31 of 48 (64.5%) cases, compared to 15 of 41 (36.6%) cases with primary tumor simplicity scores less than 0.5 (Fisher exact test p-value=0.01).

### Microenvironment transitions upon GBM recurrence

Debulking surgery, radiotherapy and chemotherapy provide therapeutic barriers but nonetheless induce tumor evolution, including influences on the tumor microenvironment. We explored this possibility by comparing the tumor associated microenvironment in primary and recurrent GBMs using CIBERSORT (**Table S8**)(Newman et al., 2015). A comparison between 91 primary and recurrent IDH-wild type tumors revealed a decrease in monocyte gene signature expression at recurrence, suggesting relative depletion of circulation derived monocytes (Figure 5A). Next, we dissected microenvironment fluctuations between diagnosis and recurrent tumors across different subtype combinations. Primary non-MES (CL or PN) tumors showed relatively high tumor purity and consequently, recurrent tumors classified as non-MES demonstrated a relatively global decrease of immune cells while cases transitioning to MES at recurrence represented a trend towards increased immune cell fractions (Figure 5B). Gene signatures of immunosuppressive regulatory T cells showed an increase in gene expression at recurrence across several primary-recurrence subtype combinations although the inferred cellular fractions are relatively small (Figure 5B). In contrast to the trend of monocyte depletion, the imputed M2 macrophage frequency was significantly higher at recurrence in cases transitioning to MES (Figure 5C). This observation converges with the higher predicted fraction of M2 macrophages in primary MES GBM relative to primary non-MES GBM. M1 macrophages and neutrophils also correlated with primary MES GBM, but these associations were not confirmed for recurrent GBM. We validated the increase in macrophages using immunostaining of IBA1 expression in two primary-recurrent GBM pairs which were classified as CL to MES (Figure 5D). IBA1 immunoexpression was restricted to macrophages/microglia, cells exhibiting either globular or filamentous/spidery morphology, with no expression in glioma tumor cells (Figure 5D). Quantitative analysis of microglia frequency using Inform software for automated pathology imaging processing confirmed a significantly higher presence (p value = 2.25e-11 and 2.12e-13 for patient #1 and #2, respectively) of at MES recurrence (Figure 5E). These findings further solidify the association between MES GBM and macrophage/microglia and extend this mutual relationship to disease recurrence. MES tumors at recurrence compared to primary MES tumors showed an increase in transcriptional activity associated with non-polarized M0 macrophages, which has been previously described (Gabrusiewicz et al., 2016), but also dendritic cells which is potentially motivated by the increased levels of neoantigens at disease recurrence (Kim et al., 2015). In contrast, primary PN GBM were found to contain significantly higher fractions of five immune cell categories compared to recurrent PN GBM, indicating a relative absence of immune infiltration in PN GBM upon recurrence.

We evaluated the effect of transcriptional class on patient survival. The analysis was restricted to 50 cases for whom annotation on overall survival (OS) time and time to disease progression (PFS) were available and with high simplicity scores, indicating low transcriptional heterogeneity. We confirmed the worse prognosis for patients whose primary tumor was classified as MES on overall survival (logrank test p=0.029 with HR=1.97)(Figure 6A; Figure 6B). This pattern was retained in patients whose secondary glioma was classified as MES (logrank test p=0.09 with HR=1.71)(Figure 6C; Figure 6D). Consequently, cases for whom both primary and recurrent tumor were classified as MES subtype showed the least favorable outcome, suggesting an additive effect of transcriptional class at different time points (Figure 6E, Figure 6F)(Figure S8).

### Treatment-Induced Immunological Microenvironment Changes upon GBM recurrence

Temozolomide treatment of gliomas can induce hypermutation(Hunter et al., 2006; Kim et al., 2015). Missense mutations may generate neoantigens that can be recognized which by CD8+ T lymphocytes (Schumacher and Schreiber, 2015). Using matching exome data we classified five recurrent gliomas in our dataset as being hypermutated at (>=400 SNVs). The predicted frequency of CD8+ T cells was significantly increased at recurrence in comparison to their primary tumors (median 7.7‰ vs 1.9‰; Wilcoxon rank test p-value=0.008)(Figure 7A). This observation was further validated by comparing seven hypermutated primary GBMs to 238 non-hypermutated GBMs (median 7.0‰ vs 0‰; Wilcoxon rank test p-value=0.031)(Figure 7B). The majority (61%) of non-hypermutated primary GBMs showed predicted CD8+ T cell fractions equal to zero. The observation suggests that patients with hypermutated tumors are more likely to benefit from CD8+ T cell antitumor immunity.

**Figure 7.**
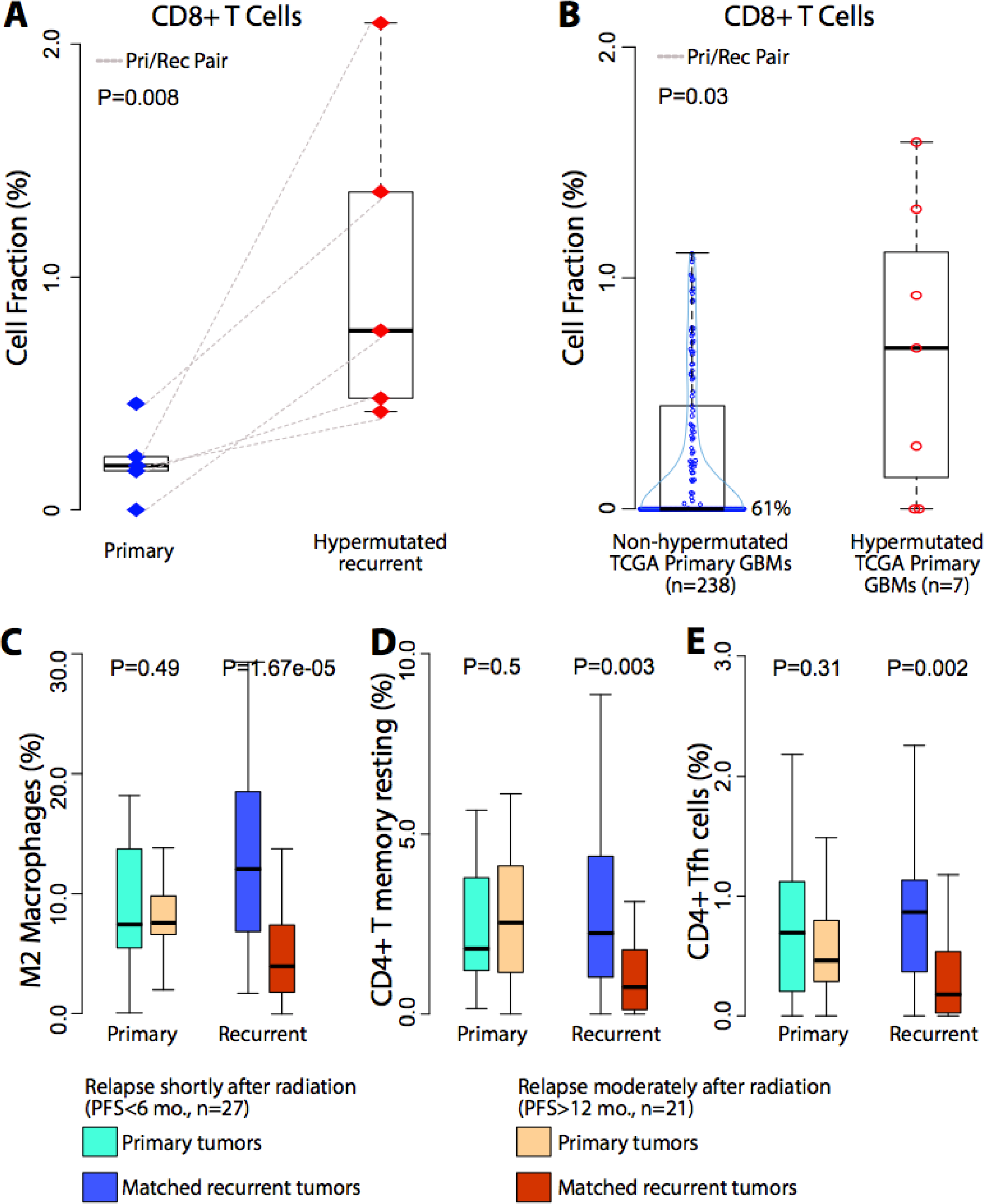
Immune cell frequency comparison. (**A**). Blue and red diamond indicate individual primary and recurrent tumors. Dash line connects paired primary and recurrent tumors. (**B**). Blue and red cycle indicate non-hypermutated and hypermutated primary samples. (**C-E**). Sky blue/dark blue and orange/red boxplots indicate short- and long- term relapsed tumors, respectively. Wilcoxon rank tests were used to examine the significance of the differences between groups.

Preclinical studies suggested radiation may increase the recruitment of T cells in the tumor microenvironment (Deng et al., 2014; Zeng et al., 2013). We compared the microenvironment of primary GBM treated with radiation therapy and separated short term relapse (PFS > 6 months, n = 27) from late relapse (PFS > 12 months, n = 21). Evaluating the gene signature based presence of M2 macrophages and CD4+ T cells (CD4+ T memory resting and CD4+ follicular helper cells) in 75 IDHwt GBM pairs whom received radiotherapy, we observed no significant difference between primary tumors with short-term and long-term relapse but found a significant increase after radiation at recurrence (Figure 7C, D, E). M2 macrophages have been speculated to play a role in resistance to radiotherapy(Meng et al., 2010; Ruffell and Coussens, 2015) and macrophage targeting immunotherapy (Pyonteck et al., 2013; Ries et al., 2014) may play a radiosensitizing role. The increasing of CD4+ T cells at recurrence for short term relapse tumors points towards inhibiting CTLA-4 as adjuvant therapy to radiation.

## DISCUSSION

Transcriptome profiling of tumor samples is a commonly used modality for interrogating pathway functionality and phenotype based patient classification. The transcriptional footprint left by the tumor microenvironment, which may constitute 10-80% of cells in a tumor biopsy(Yoshihara et al., 2013), can obscure the true activity of the signaling network(Isella et al., 2015; Kim and Verhaak, 2015). Here, we employed in silico methods to integrate mRNA expression profiles from glioma samples and glioma cell culture models to provide insights into glioma-intrinsic pathway activities and classification, and to deconvolute the glioma associated stroma into its immunological cellular components.

GBM expression subtype classification has emerged as an important concept to better understand the biology of this devastating disease (Dunn et al., 2012; Huse et al., 2011; Sturm et al., 2014). Robust classification of new GBM tumors is therefore critical to ensure consistency in reporting between different studies. The transcriptional glioma subtypes we discovered using tumor-intrinsic gene expression values strongly overlapped with the proneural, classical and mesenchymal subtypes but identified the neural class as normal neural lineage contamination. Our updated methods, released through a R-library, were found to be highly robust and provide the community with a standardized strategy for classification of gliomas.

Through re-classification of primary GBM samples from TCGA and despite using tumor-only transcripts, we observed that the mesenchymal GBM subtype associated with the presence of tumor-associated glial and microglial cells. Mesenchymal glioma cell differentiation status has been found to correlate with enrichment of macrophages/microglia (Bhat et al., 2013; Kreutzberg, 1996). Through in silico cell type identification we additionally detected enrichment of various adaptive immunity cell types, including CD4+ T lymphocytes.

Longitudinal analysis of tumor samples is complicated by the lack of tissue collections including such pairs. Through aggregation of existing and novel datasets we compiled a cohort of 124 glioma pairs, including 91 pairs of IDH wild type tumors. Comparison of pairs of initial gliomas and first disease recurrence did not identify the trend of proneural GBM transitioning to a mesenchymal phenotype that has often been suspected (Phillips et al., 2006). Mesenchymal subtype at diagnosis and at disease recurrent correlated with relatively poor outcome. The recurrent IDH wild type GBM immuno environment showed fewer blood derived monocytes which may reflect lower penetration through the blood brain barrier. While the frequency of M2 macrophage/microglia was increased in recurrent mesenchymal GBM compared to primary non-mesenchymal GBM, the overall fraction of M2 macropage/microglia remained stable. This possibly suggests that the majority of these cells are derived from resident CNS macrophages than through active recruitment from the circulation.

In summary, our study defines a new strategy to determine transcriptional subtype, and associated expression classes to the tumor-associated immuno-environment. Our findings may aid in the implementation of immunotherapy approaches (Blank et al., 2016) in a disease type with very limited treatment options. Collectively, our results have improved our understanding of determinants of GBM subtype classification, the critical impact of the tumor microenvironment, and provide new handles on the interpretation of transcriptional profiling of glioma.

## EXPERIMENTAL PROCEDURES

### Collection of pairs of primary and recurrent glioma samples

U133A array profiles for 543 primary GBM, and RNA-Seq data for 166 primary and 13 recurrent GBM were obtained from the TCGA portal https://tcga-data.nci.nih.gov/tcga/. Mutation calls and DNA copy number profiles were obtained for all samples, where available. Tissues from 20 additional initial GBM and matched recurrent tumor were obtained from Henry Ford Hospital (n = 9) in accordance with institutional policies and all patients provided written consent, with approval from the Institutional Review Boards (Henry Ford Hospital IRB protocol #402). All RNA samples tested were obtained from frozen specimens. All of the recurrent GBMs had been previously treated with chemotherapy and radiation. Three cases had a history of lower grade astrocytoma prior to the first GBM (HF-2869/HF-3081/HF-3162). Tumors were selected solely on the basis of availability. RNA-Seq libraries were generated using RNA Truseq reagents (Illumina, San Diego, CA, USA) and paired-end sequenced using standard Illumina protocols. Read length was 76 base pairs for cases sequenced by TCGA and from Henry Ford (processed at MD Anderson). RNA-Seq data on frozen tissue from 44 patients with initial and recurrent GBM that received resection at Samsung Medical Center and Seoul National University Hospital were provided by Dr. Nam’s lab. Surgery specimens were obtained in accordance to the Institutional Review Board (IRB) of the Samsung Medical Center (No. 2010-04-004) and Seoul National University Hospital (No. C-1404-056-572). Affymetrix CEL files of 39 pairs of initial and recurrent glioma were retrieved from the Gene Expression Omnibus (GEO accession GSE4271, GSE42670, GSE62153)(Joo et al., 2013; Kwon et al., 2015; Phillips et al., 2006). The expression profiles of the 23 pairs from GSE4271 were determined using Affymetrix HG-U133 GeneChips, the 1 pairs from GSE42670 were analyzed using the Affmetrix HuGene-1-0-st platform, the 15 pairs from GSE62153 were analyzed using Illumina Human HT-12 V4.0 expression BeadChip. The RNA sequencing data of 14 and 5 pairs of primary and recurrent low grade glioma were from TCGA LGG cohort and EGAS00001001255(Mazor et al., 2015), respectively.

Genome wide DNA copy number profiling and exome sequencing on thirteen TCGA tumor pairs and nine of ten Henry Ford tumor pairs were performed and data was analyzed using standard protocols and pipelines as previously described (Kim et al., 2015).

### Data for multiplatform classification comparison

RNA sequencing data was available for 162 primary GBMs(Brennan et al., 2013) for which an Affymetrix HT-U133A gene expression profile was also available. We observed a low Pearson Correlation Coefficient (< 0.15) between RNA sequencing based reads per kilo base of transcript per million reads (RPKM) and Affymetrix HTU133A profiles in eighteen cases and these were removed from further analysis. In summary, in order to assess the concordance between classification results of the new 70-gene signatures and previously published 210-gene signatures (Verhaak et al., 2010b), 144 GBMs which were profiled in both RNA sequencing and Affymetrix U133A platforms were used in our further analyses.

### Transcriptome Data processing

The latest version custom CDF files (Version19, http://brainarray.mbni.med.umich.edu) (Dai et al., 2005; Sandberg and Larsson, 2007) were used to map probes from the Affymetrix HG-U133A and HuGene-1_0-st GeneChip platforms to the Ensemble transcript database, combined in one probe set per gene and normalized using the AROMA package with default parameters, resulting in RMA normalized and log transformed gene expression values (Bengtsson et al., 2009). All RNA sequencing data was processed by the PRADA pipeline (Torres-Garcia et al., 2014). Briefly, reads were aligned using BWA against the genome and transcriptome. After initial mapping, the aligned reads were filtered out if their best placements are only mapped to unique genomic coordinates. Quality scores are recalibrated using the Genome Analysis Toolkit (GATK), and duplicate reads are flagged using Picard. Mapped features were quantified and normalized per kilo base of transcript per million reads (RPKM) and were converted to a log2 scale to represent a gene expression level. RPKM values measuring the same gene that mapped to the Ensemble transcript with longest size were selected to obtain one expression value per gene and sample. RPKM values were converted to a log2 scale to represent gene expression level. The statistical environment R was used to perform all the statistical analysis and graph plots.

### Deriving new gene signatures

A pair-wise gene expression analysis identified 5,334 genes which are significant higher expressed in glioma bulk samples compared to their derivative GSCs. These genes were discarded from the gene list for developing tumor-specific molecular subtypes. Consensus non-negative matrix factorization (CNMF) clustering method identified three distinct subgroups among the 369 IDHwt primary GBMs. A set of 270 GBMs was recognized as core samples based on a positive silhouette width. The gene expression values of each subtype were compared with those from the other two subtypes combined (Verhaak et al., 2010b). Signature genes per cluster were selected on the basis of differences in gene expression level and were considered significant if they reached the cut-off value with t-test p-value< 1E-3 for higher expressed in this class, while also showing a significant lower expression with t-test p-value<1E-3 in the other two classes. In the original gene signatures, genes could be either down-regulated or up-regulated, while only up-regulated genes (n=70 per gene signature) were selected for revised gene signatures. Only genes measured on both RNAseq and U133A platforms were considered, and the U133A data from 162 GBM samples measured on both platforms (which included the 144 cases used to compare U133A and RNAseq results) was used in the final comparative analysis.

### Molecular classifications based on ssGSEA enrichment scores

Single sample gene set enrichment analysis was performed as follows. For a given GBM sample, gene expression values were rank-normalized and rank-ordered. The Empirical Cumulative Distribution Functions (ECDF) of the signature genes and the remaining genes were calculated. A statistic was calculated by integration of the difference between the ECDFs, which is similar to the one used in GSEA but is based on absolute expression rather than differential expression (Barbie et al., 2009). Since the ssGSEA test is based on the ranking of genes by expression level, the uncentered and log-transformed U133A and RPKM expression levels were used as input for ssGSEA. Since the scores of the three signatures were not directly comparable, we performed a resampling procedure to generate null distributions for each of the four subtypes. First we generated a large number of virtual samples in which each gene obtains its expression level by randomly selecting an expression value of the same gene in the remainder of the samples. Then, the three ssGSEA scores for each signature were calculated. Following this procedure we generated a large number (>1,000,000) of random ssGSEA scores for each subtype, to build the null distribution and to give empirical p-values for the raw ssGSEA scores of each sample. By testing on multiple datasets with different sample sizes, we found the resampling generated distribution could be replaced with Student-T distribution (sample size>30) or Normal distribution (sample size>50) for getting very similar results. R-library with the code and expression matrices used is provided as supplementary file..

### Evaluate the simplicity of subtype activation

For a single sample, we decreased rank the empirical p-values for each subtype to generate order statistics as *R*_*N*–1_, *R*_*N*–2_ … *R*_1_, *R*_0_. In particular, *R*_0_ equals to the minimum empirical p-value and points to the dominant subtype, i.e., the most significantly activated subtype. The accumulative distance to the dominant subtype (ADDS) was defined as:

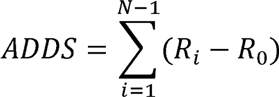

Similarly, the accumulative distance between non-dominant subtypes (ADNS) as:

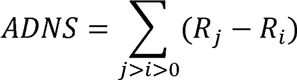

Obviously, the ADDS and ADNS are positive and negative correlated with single activation, respectively. Hence, we defined the simplicity score by combing ADDS and ADNS together and corrected with a constant 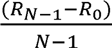 as follows:

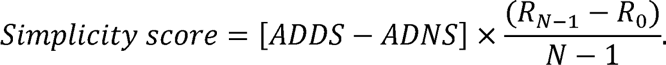

### Tumor purity assessment

The ESTIMATE package was used to evaluate tumor purity on the basis of the expression level of marker genes in stromal and immune cells (Yoshihara et al., 2013), where the fraction of stromal cells and immune cells in each sample were represented by stromal score and immune score respectively, and the mixed fraction of both stromal and immune cells was represented by estimate scores. The ABSOLUTE package was used to confirm the tumor purity on the basis of chromosome copy number and allele fraction ratios on samples for which single nucleotide polymorphism array data were available (Carter et al., 2012).

### Sample collection and Neurosphere Cultures

After obtaining approval from the institutional review board of The University of Texas M.D. Anderson Cancer Center, glioblastoma tumor tissues were collected and named in the order that they were acquired. Each tissue was enzymatically and mechanically dissociated into single cells and grown in DMEM/F12 media supplemented with B27 (Invitrogen), EGF (20 ng/ml), and bFGF (20 ng/ml), resulting in neurosphere growth. All cell lines were tested to exclude the presence of Mycoplasma infection. To minimize any batch effect the downstream molecular analyses were performed on identical cell culture batches. Total RNA from formalin fixed, paraffin embedded tumor tissues and matching neurospheres was prepared using the Masterpure complete DNA and RNA isolation kit (Epicenter) after proteinase K digestion per to the instructions from the manufacturer. Paired-end Illumina HiSeq sequencing assays were performed resulting in a medium number of 50 million 75bp paired end reads per sample. We employed the PRADA pipeline to process the RNA sequencing data (Torres-Garcia et al., 2014). In short, Burroughs-Wheeler alignment, Samtools, and Genome Analysis Toolkit were used to map short reads to the human genome (hg19) and transcriptome (Ensembl 64) and RPKM gene expression values were generated for each of the 135,994 transcripts of 21,165 protein coding genes in Ensembl database.

### Western blotting

Lysates were prepared from fresh frozen sections using RPPA lysis buffer (1% Triton X-100 50mM HEPES pH 7.4, 150mM NaCl, 1.5mM MgCl2, 1mM EGTA, 100mM NaF, 10mM Na pyrophosphate, 1mM Na3VO4, 10% glycerol, plus protease and phosphatase inhibitors cocktails from Roche Applied Science #05056489001 and 04906837001), with sonication and clearing by centrifugation at 10,000g. Protein concentration was measured using the BCA kit (Thermo Scientific - Pierce #23225). SDS-PAGE and western blotting was performed using Midi gel system (Life Technologies - #WR0100) and NuPage-Novex 4-12% Bis-Tris Midi (20-well) Protein Gels (Life Technologies - #WG1402) using the following antibodies: ITGAM (CD11B) (Sigma Aldrich – #HPA002274), IBA1 (AIF1) (Sigma Aldrich – #HPA049234), GFAP (Cell Signalling – #3670), YKL40 (CHI3L1) YKL40 (CHI3L1, Santa Cruz Biotechnology - #sc-30465) a-actinin (Sigma Aldrich A5044) and Tubulin (Sigma Aldrich T9026).

### Immunohistochemistry

Formalin-fixed, paraffin-embedded tissue sections (4 µm thick) were collected on Superfrost plus slides. Briefly, tissue sections were deparaffinized with xylene and ethanol and re-hydrated with 95, 70 and 50% ethanol. Sections were antigen unmasked using citrate buffer (Vector Labs #H-3300) and heating. Peroxidase block was conducted with 3% H2O2 and blocking was with 5% goat serum (Vector Labs #S-1000). Primary rabbit polyconal antibody against IBA1 (AIF1)(WAKO #016-20001) at 1:400 was used overnight. Secondary antibody was done using with the Rabbit-on-Rodent HRP-Polymer (Biocare #RMR622L) for 1 hr at room temperature. The slides were developed with Nova-red (Vector Labs #SK-4800) and counterstained with haematoxylin, mounted and scanned with Pannoramic 250 slide scanner (Caliper Life Sciences). Unbiased quantification of microglial (IBA1+) percentage in primary and recurrent GBMs was performed using the Caliper Vectra image system and InForm analysis software. Thirty scan fields were automatically selected on from entire tumor section. Nineteen scan fields were select from the primary tumor of patient #2 due to the small size of tumor section. Percentages of the median and high levels (2+, 3+) of IBA1 were used for the comparison.

## Acknowledgments

The authors thank Katherine Stemke-Hale for assistance in manuscript editing. The results published here are in part based upon data generated by The Cancer Genome Atlas project established by the National Cancer Institute (NCI) and the National Human Genome Research Institute (NHGRI) of the National Institutes of Health. Information about TCGA and the investigators and institutions that constitute the TCGA research network can be found at http://cancergenome.nih.gov. This work is supported by grants from the National Institutes of Health grants P50 CA127001 (EPS, RGWV), R01 CA190121 (EPS, RGWV), P01 CA085878 (RGWV); RO1 CA120813 (ABH), and P30 CA016672 (MD Anderson Cancer Center Support Grant for the Sequencing and Microarray Facility); Cancer Prevention & Research Institute of Texas (CPRIT) grant number R140606, the University Cancer Foundation via the Institutional Research Grant program at the University of Texas MD Anderson Cancer Center (RGWV); the National Brain Tumor Association Defeat GBM project (EPS, RGWV), the National Brain Tumor Association Oligo Research Fund (RGWV); the Korea Health Technology R&D Project through the Korea Health Industry Development Institute (KHIDI), funded by the Ministry of Health & Welfare, Republic of Korea (HI14C3418, DHN).

## Supplementary Information

### Supplementary Figures and Legends

**Figure S1.**
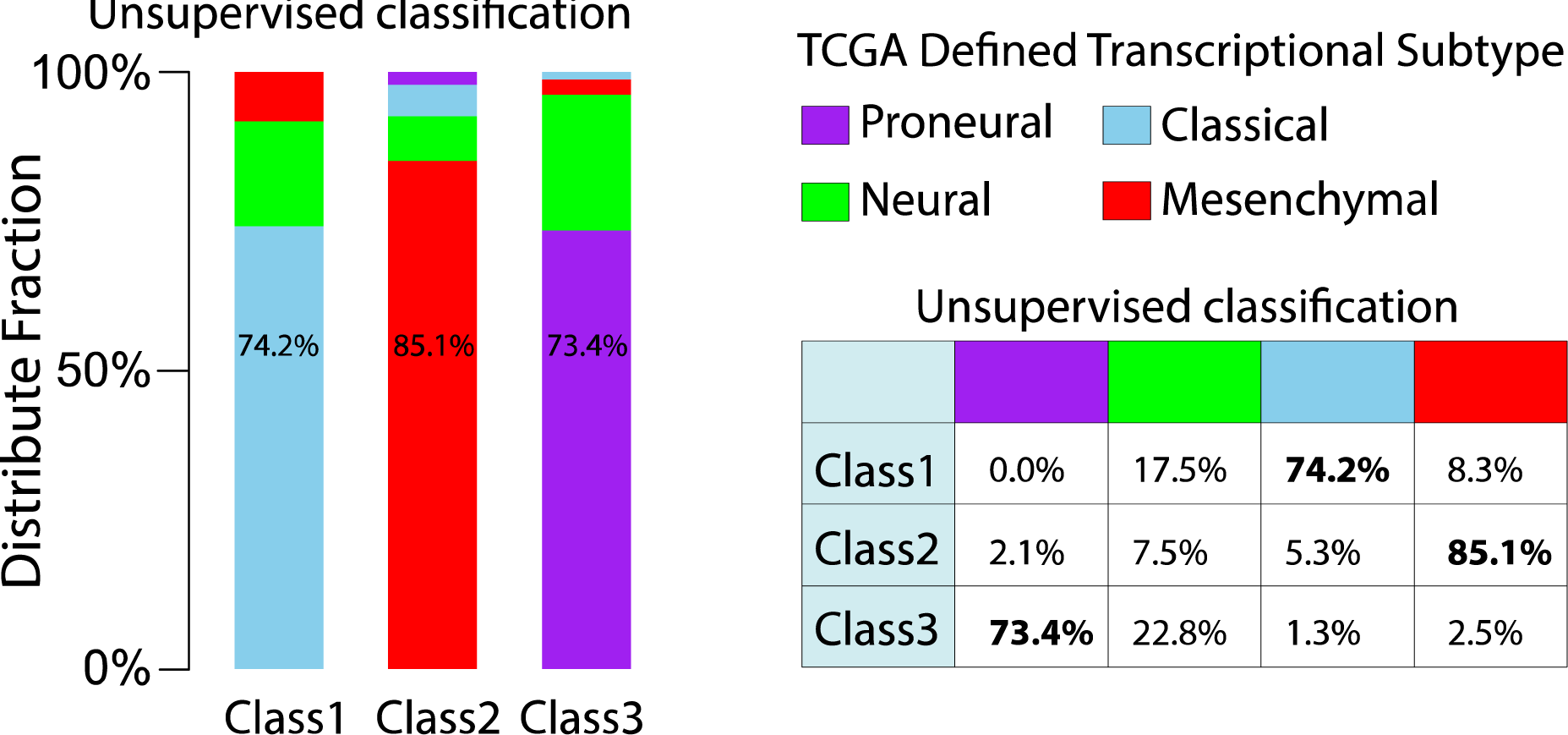
Comparison between GCIMP-GBM specific classification and TCGA defined GBM subtypes. 270 samples were identified as core samples with positive silhouette width core samples. 97, 94 and 79 samples were unsupervised classified class1, class2 and class3, respectively. The previous four transcriptional subtypes of these 270 samples were determined by TCGA Research Network (Brennan et al., 2013).

**Figure S2.**
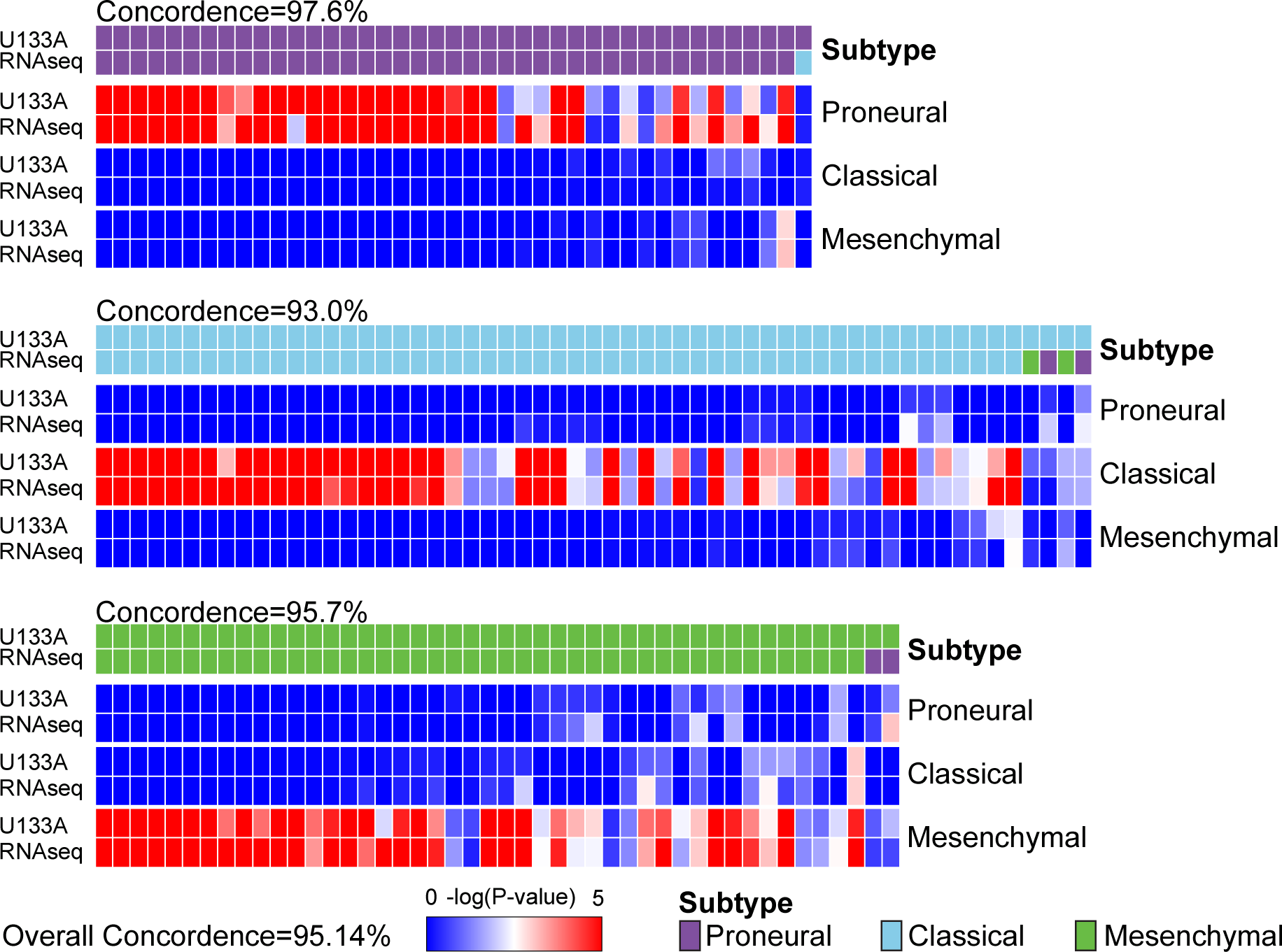
Concordance of transcriptional classification of GBMs crosses multiple platforms. Through TCGA, the expression profiles of 144 GBM were analyzed using both Affymetrix U133A gene expression arrays and RNA sequencing. The empirical -log(P-value) of raw ssGSEA enrichment scores at each signature are shown as heatmaps, with dark blue representing no activation and bright red as highly activated. For each panel, the first row shows U133A based classification, and the second row indicates RNA-seq subtype classification.

**Figure S3.**
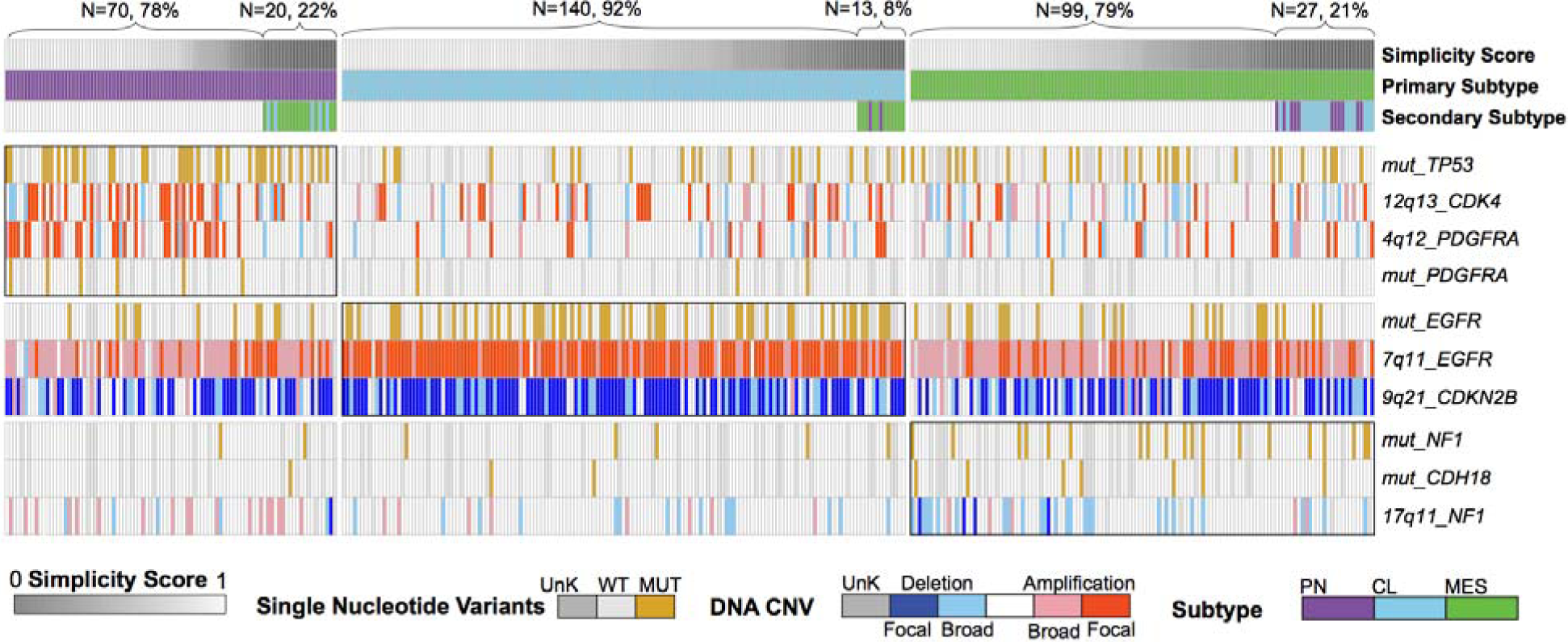
Genomic alteration patterns for each subtype. Ten prominent GBM somatic events were manually selected. Simplicity score and primary subtype were shown by the first two rows. For samples with simplicity score less than 0.1, the secondary subtype was also shown by the third row.

**Figure S4.**
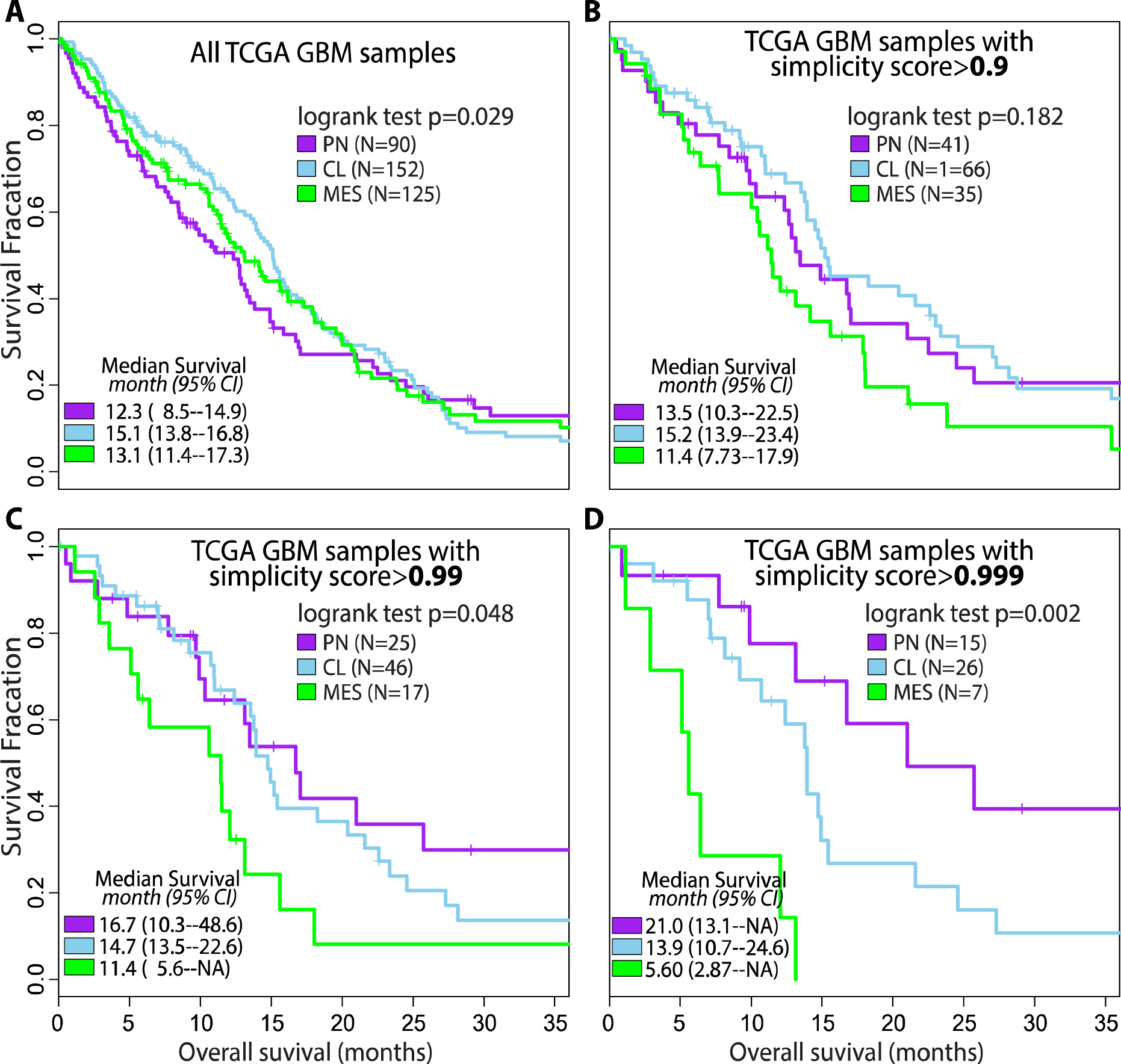
Patient survival differences between transcriptional subtypes. Samples were filtered using increased simplicity score as threshold from panel A to D.

**Figure S5.**
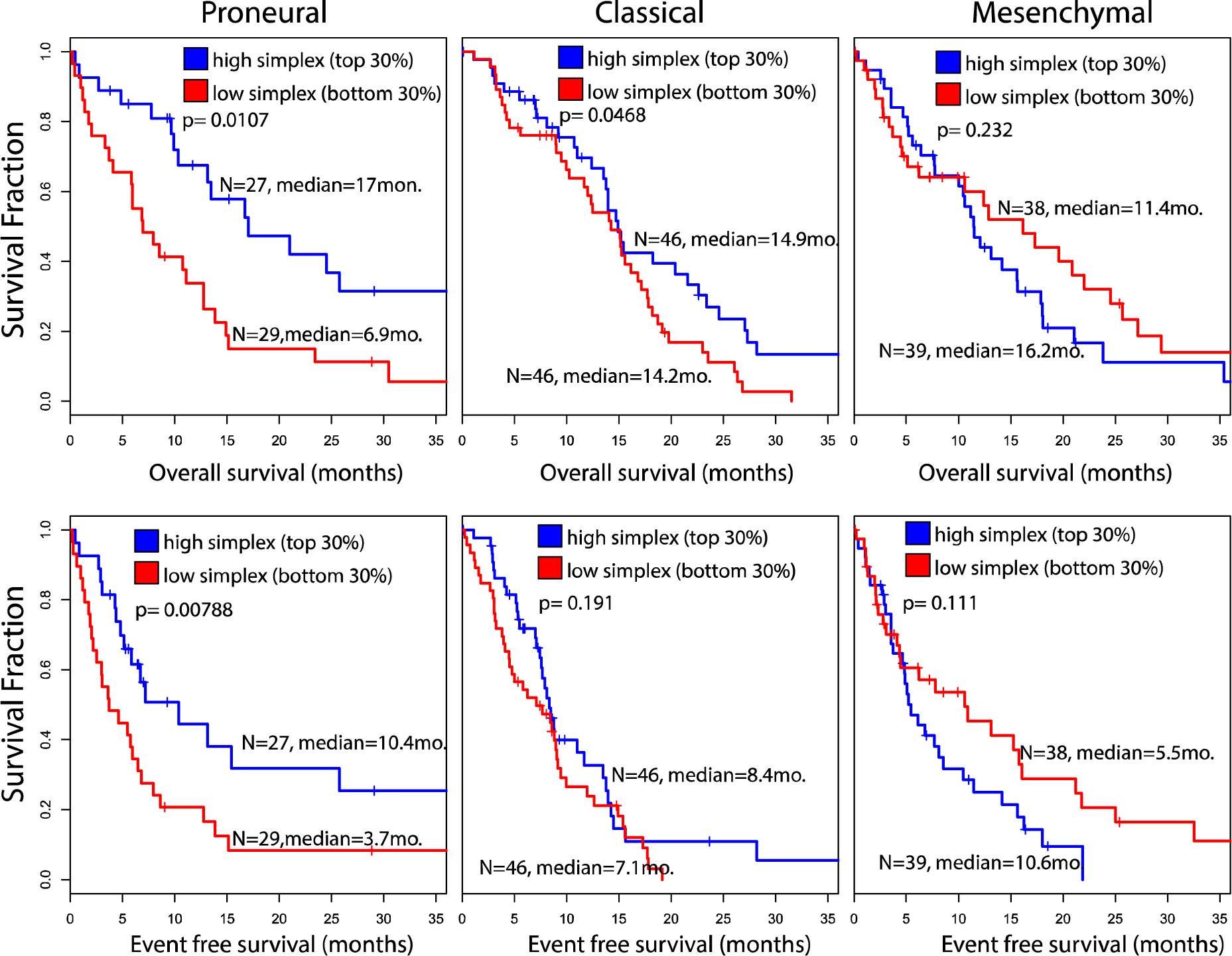
Overall and event free survival analysis comparison between samples with high and low simplicity scores in each subtype.

**Figure S6.**
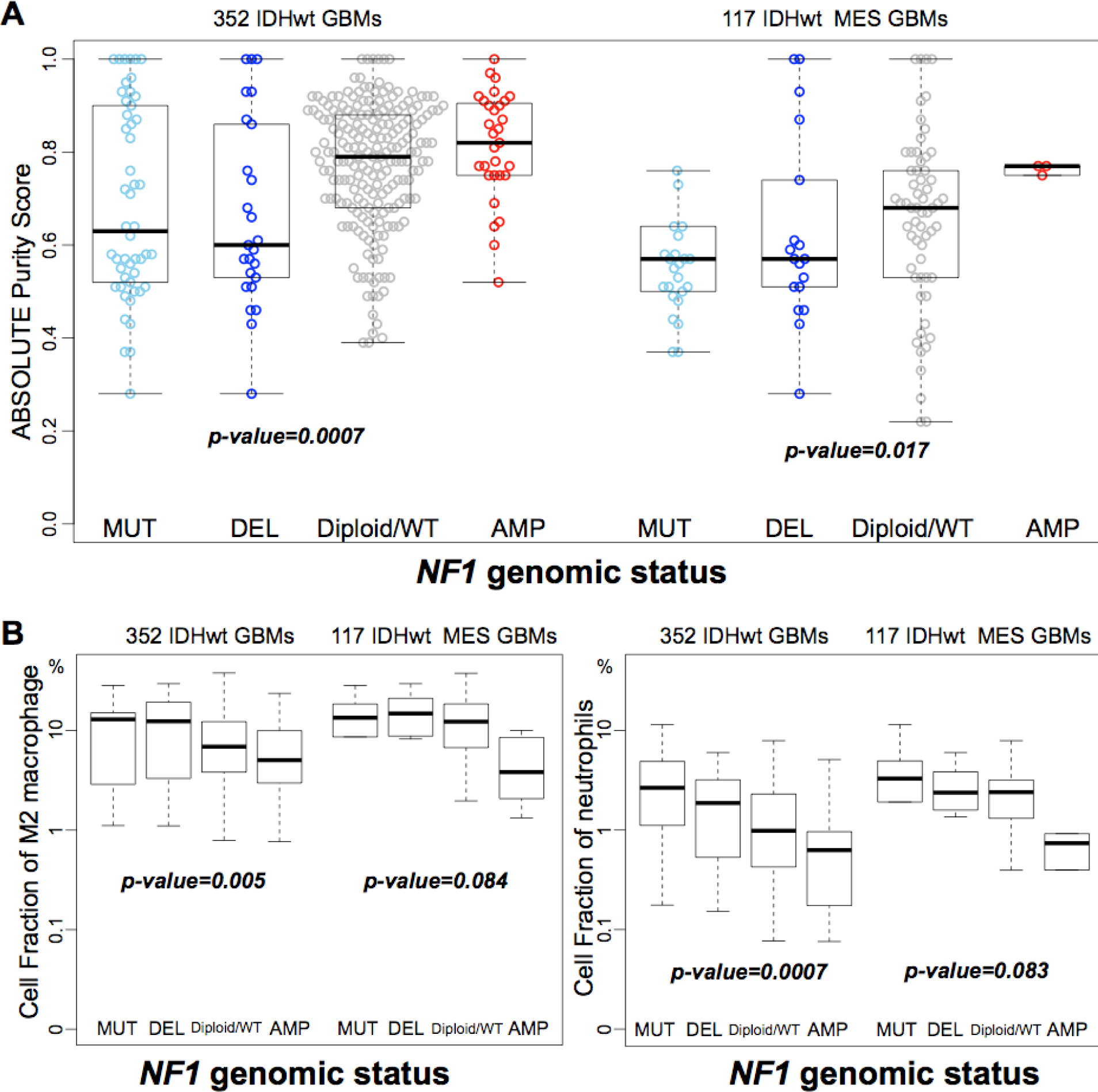
Comparison of tumor purity and immune cell fraction between GBMs with different *NF1* genomic status. P-values were calculated using Wilcoxon rank test between samples carrying *NF1* deletion/mutation and others.

**Figure S7.**
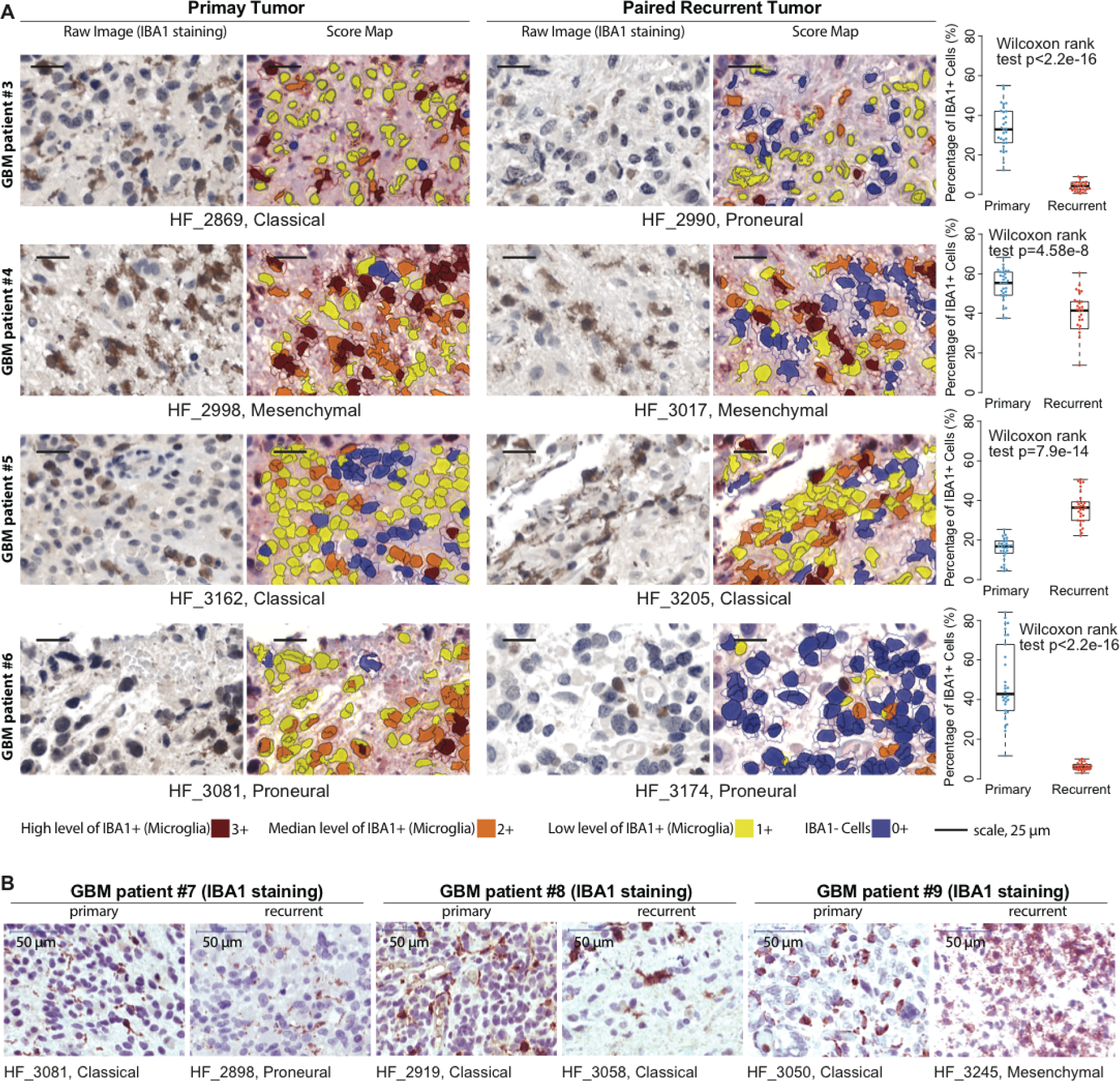
Immunohistochemistry staining of the IBA1 imcroglia marker. (**A**) Representative images with immunohistochemical staining of the IBA1 and score map obtained by InForm image analysis tools in four matched pairs of primary and recurrent GBM. Thirty scan fields were unbiased selected for each tumor by Calipar Vectra pathology imaging system automatically. (**B**) IHC staining of the IBA1 in three additional matched pairs of primary and recurrent GBM.

**Figure S8.**
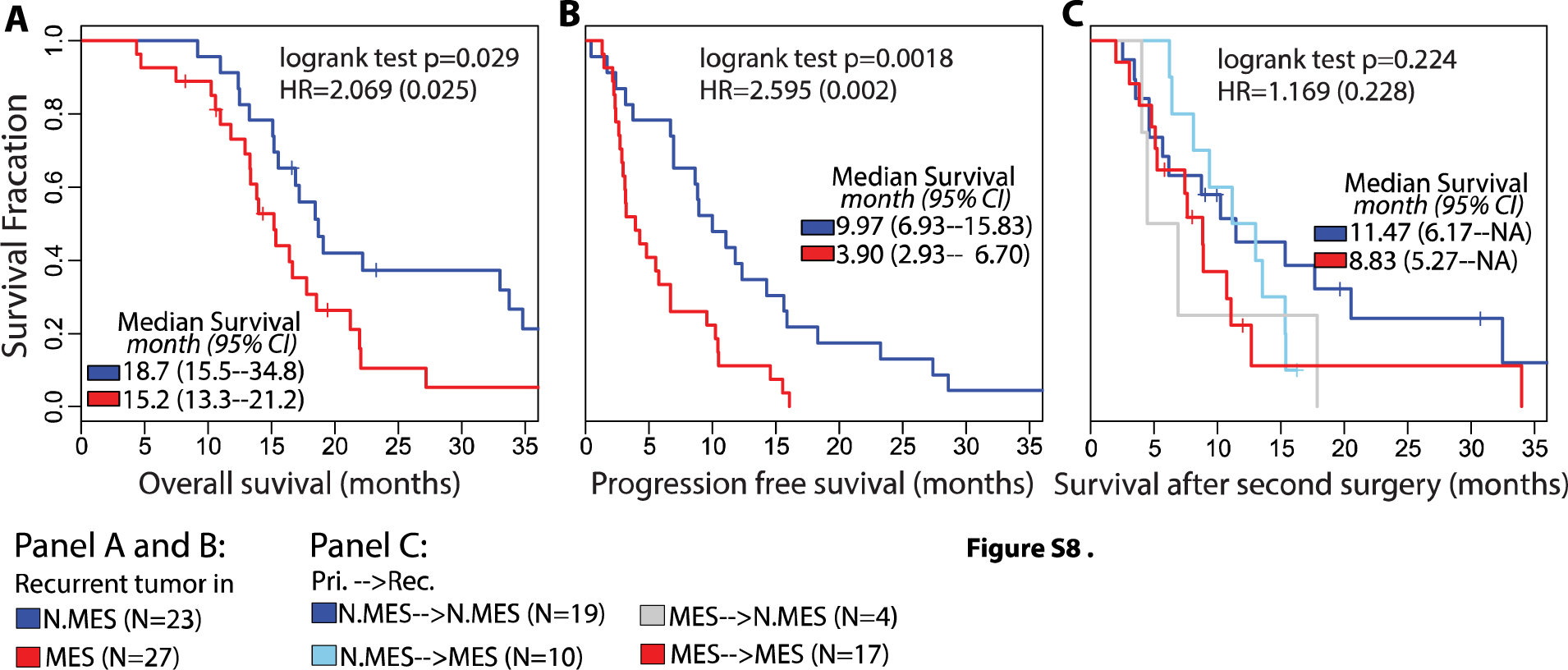
**(A, B)** Overall and progression free survival analysis between samples in different recurrent subtypes. **(C)** Survival after secondary surgery comparison between different transition types.

